# EEG responses to auditory cues predict fluency variability and stuttering intervention outcome

**DOI:** 10.1101/2025.02.21.635719

**Authors:** Monica Filipa Rocha, Jaqueline Carmona, João Mendonça Correia

**Author notes:** these authors participated equally for the study) (Monica Filipa Rocha, Jaqueline Carmona).

## Abstract

Stuttering is a variable speech disorder whose brain mechanisms remain unknown. Sensorimotor brain circuits, critical for motor-speech control, including auditory processing necessary for speech prediction and monitoring, have been linked to the disorder. Despite considerable advances, it remains unclear whether auditory circuits relate to stuttering variability, and whether the panoply of interventions for persons who stutter can lead to brain changes within these circuits. We employed electroencephalography (EEG), in a group of persons who stutter, in combination with auditory probes to tap onto the importance of auditory cortical regions in stuttering variability. Participantsproduced flexible speech (i.e., describing visual scenes) and non-flexible speech (i.e., reading syllables), following an auditory cue. More pronounced P200 auditory evoked potentials were observed in participants with higher dysfluency rates, mainly in the spontaneous speech task. Interestingly, speech therapy intervention led to a reduction of the P200 potential, which was in turn significantly related to fluency improvements. Furthermore, EEG response patterns discriminative of cue frequency (400 or 800 Hz tones) were also predictive of dysfluency scores. Our study highlights the involvement of auditory cortical processing and that of auditory attention in stuttering variability. We support that a higher state of auditory alertness may be implicated in the sensorimotor mechanisms of stuttering, and that speech therapy interventions promoting more self-confident communication can restraint auditory alertness, and potentially reduce speech dysfluencies.

**Highlights:**

1. Auditory probes can assess the auditory cortex in speech production and stuttering.
2. Stuttering severity correlates to EEG auditory responses during speech preparation.
3. Higher states of auditory alertness in stuttering may be reduced by speech therapy.

**Graphical abstract:** 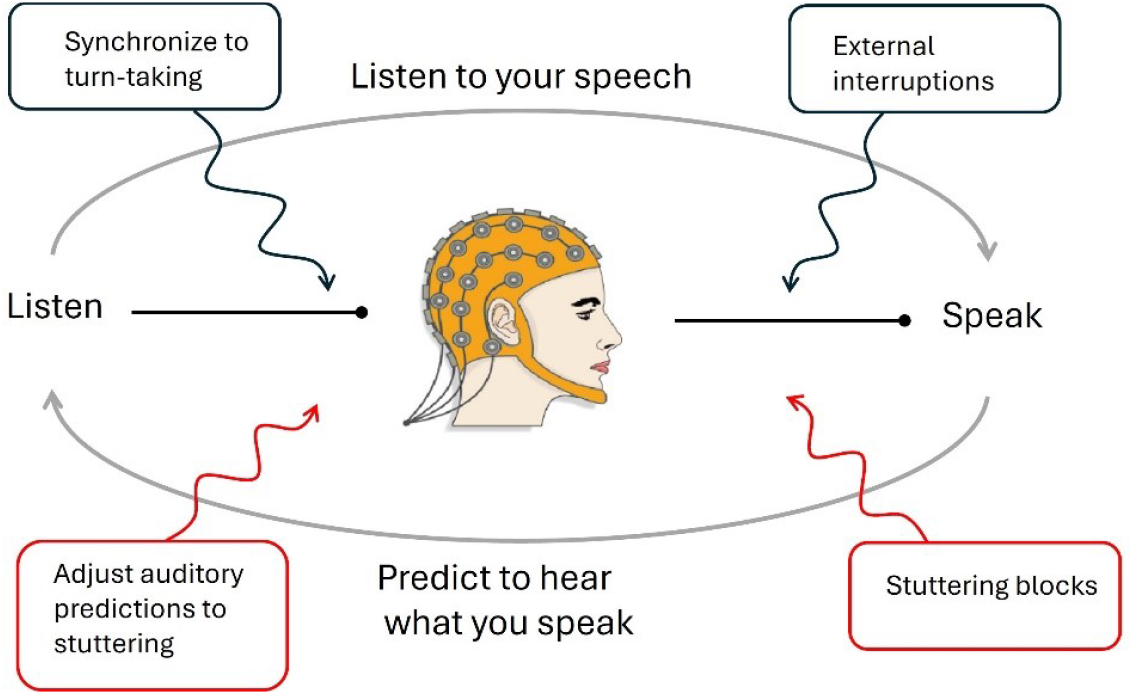

Speaking requires orchestrating several brain processes at a time. The auditory system assumes a central role, not only in waiting for the right moment to initiate speech, listening to self-produced speech, predicting the consequence of future speech, but also adjusting these processes to the intermittent nature of stuttering.

## 1. Introduction

Persistent developmental stuttering (PDS) is a communication disorder that impacts 1% of the world population (Yairi and Ambrose, 2013). To date, its neural mechanisms remain unknown, limiting our capacity to develop effective brain-informed strategies to curb stuttering. Worldwide, stuttering recovery is age dependent, with a remarkable high rate in children over 80% (Dworzynski *et al*., 2007; Yairi and Ambrose, 2013) and an odd low rate in adults, suggesting that neural networks responsible for speech development may be implicated in the disorder (Chang and Zhu, 2013; Neef, Anwander and Friederici, 2015). Neuroimaging studies indicate that sensorimotor circuits required for speech development are also involved in everyday speech control (Poeppel *et al*., 2012; Dick, Bernal and Tremblay, 2014; Tremblay, Deschamps and Gracco, 2016), including prediction and monitoring of the sensory consequences of speech production (i.e., auditory and somatosensory) (Ohashi and Ostry, 2021). Attenuation of brain activity in auditory cortical regions, within the superior temporal lobe, is generally observed in response to self-generated speech sounds, which may reflect motor-to-auditory prediction of auditory speech consequences (Crapse and Sommer, 2008; Christoffels *et al*., 2011). Persons who stutter (PWS) show altered auditory attenuation in comparison to persons who do not stutter (PWNS) (Brown *et al*., 2005; Beal *et al*., 2010). Additionally, auditory-to-motor cortical connections are suggested to be involved in speech perception (Correia, Jansma and Bonte, 2015) and in correcting on-going speech gestures to match auditory targets (Guenther, Ghosh and Tourville, 2006; Hickok, Houde and Rong, 2011). When acoustic modulations are imposed to auditory feedback, the motor system quickly adapts the articulatory targets to reduce the mismatch between predicted and perceived speech feedback (Kalinowski *et al*., 1993). Furthermore, it has been shown that in certain altered auditory feedback (AAF) conditions, PWS have slower motor adaption in comparison to PWNS (Cai *et al*., 2012). Interestingly, PWS tend to improve speaking fluency under AAF, including delay auditory feedback (DAF), frequency auditory feedback (FAF) or amplitude auditory feedback (AAF) (Lincoln, Packman and Onslow, 2006; Fiorin *et al*., 2021). Moreover, speaking conditions that drive attention to external auditory signals, such as choral speaking, metronome-paced speaking, shadowing speech, theatrical speech or singing, have been widely recognized as conditions that can enhance fluency in PWS (Toyomura, Fujii and Kuriki, 2015). Overall, the relationship between auditory processing and stuttering is paramount to the disorder, with several fluency enhancing techniques exploring the phenomenon (Sommer *et al*., 2002).

Beyond cortical sensorimotor circuits, the basal ganglia (BG) may be particularly implicated in stuttering (Alm, 2004; Giraud *et al*., 2008; Kubikova *et al*., 2014; Chang and Guenther, 2020). Stuttering shows several parallels to Parkinson’s disease (PD), whose neural anatomical locus lays within the BG, and whose symptoms include difficulties to initiate and stop motor actions (Alm, 2004; Toft and Dietrichs, 2011). Like stuttering, motor execution in PD can be facilitated when triggered by external cues (Sares *et al*., 2019). The direct pathway (DP) and the indirect pathway (IP) have been at the centre of BG accounts for both, PD and stuttering (Metzger *et al*., 2018). However, more recently, the hyperdirect pathway (HDP) has been investigated as a critical BG mechanism for motor control, resetting (i.e., preparing) motor cortical regions prior to motor action (Jorge *et al*., 2022; Nambu *et al*., 2023). We previously found that phonation, but not articulation, was decodable from BG nuclei during speech production (Correia *et al*., 2020) by comparing overt versus whispered speech. It is possible that the auditory cortex and the BG interact in speech production, and that an imbalance in auditory prediction and monitoring of speech consequences affect the internal rhythm by which the BG promotes fluent speech (Chang and Zhu, 2013). In which way auditory processing of self-generated speech impacts the basal ganglia circuits necessary for timely motor sequences, remains unknown.

In neuroimaging research, evaluating the role of auditory processing during speech production has revealed challenging due to concomitant signal artifacts generated by head motion (e.g., susceptibility artifacts in MRI) (Gracco, Tremblay and Pike, 2005) and respiratory confounds (Chang and Glover, 2009; Power *et al*., 2017) and electrical activity of speech muscles (e.g., interference of EEG/MEG signals) (Ganushchak, Christoffels and Schiller, 2011). Nevertheless, an interesting strategy previously employed in EEG, made use of auditory probes presented during speech preparation of individual words (Daliri and Max, 2015, 2018). Using a neuroimaging technology with high temporal resolution (e.g., EEG) and an epoch of interest that avoids overlap with speech production proper, authors reduce contamination of speaking artifacts in the brain signal. Differences in auditory evoked potentials (e.g., the P200 potential) between PWS and PWNS, during speech planning and motor preparation, suggest that the auditory cortex is critically involved in speech production and stuttering. However, while PWS may encounter difficulties in the production of isolated items, such as words, it is in spontaneous and continuous speech that stuttering has a marked impact in their speaking capabilities (Tsiamtsiouris and Cairns, 2013; Warner *et al*., 2023). Employing speech production tasks that involve varying levels of linguistic complexity, from syllables to spontaneous conversation, can provide valuable insights into the specific challenges faced by PWS (Sengupta *et al*., 2017).

Moreover, speech therapy interventions have made substantial improvements over the past decades (Brignell *et al*., 2020; Laiho *et al*., 2022). Stuttering modification techniques, based on strategies to teach PWS to adopt a softer speaking method have been replaced by methods that focus instead on the goal of communication, develop a positive self-perception of stuttering, acceptance of fluency limitations in the context of fluency diversity, and the respect for the individual speaking style (Bloodstein and Ratner, 2008). Beyond fluency, stuttering affects communication confidence (Klompas and Ross, 2004; Craig, Blumgart and Tran, 2009; Koedoot *et al*., 2011). By targeting these central aspects of stuttering, but often perceived as secondary, speech therapists tackle foundational characteristics of a PWS, which is related to speaking avoidance and life-style constraints due to stuttering (Sønsterud *et al*., 2020). Self-perception questionnaires are crucial in evaluating and treating stuttering, as they delve into how individuals perceive and experience their stuttering (Yaruss, 2010). These tools are designed to assess the subjective experiences and attitudes of PWS, providing valuable insights into the psychological and social aspects of stuttering.

In this study, we combine a modern stuttering intervention approach with electrophysiology to investigate the relation between auditory evoked potentials obtained from auditory cues during speech preparation and individual stuttering variability. Adults who stutter were volunteers in two EEG sessions, scheduled before and after a 12-week intervention program for stuttering. In the EEG sessions, participants performed a series of speech tasks combined with EEG recordings, including the production of syllables and description of images (see figure 1). We then assessed the relationship between fluency and EEG variability concerning the processing of auditory cues probing the auditory brain regions during speech preparation. Importantly, this approach yields EEG data free from articulatory EEG artifacts. Although indirect, EEG responses to auditory cues offer an indication of the involvement of the auditory cortex in speech preparation, namely the prediction of the auditory consequences of speech, a key sensorimotor aspect in speech production and stuttering. As such, we hypothesize that EEG responses to the auditory cues may vary in relation to stuttering and fluency variability, as well as in relation to intervention outcome. Furthermore, auditory cues were two tones with different acoustic frequencies (i.e., 400 and 800 Hz). Beyond investigating EEG activity time-locked to auditory cue processing, tone variation enabled the possibility to assess tonal discrimination in the EEG responses, by exploiting multivariate pattern analysis (Hausfeld *et al*., 2012). Whereas the tone presentation time is relevant for the speech task, because it serves as the cue required to initiate speech, tonal discrimination is not explicitly part of the participants’ task. Hence, EEG-based tonal discrimination can probe a more nuanced aspect of auditory processing that may contribute to our understanding of how sensory inputs influence speech motor control in individuals who stutter, potentially revealing differences in neural sensitivity or processing efficiency that are not directly observable through behaviour measures and activity-based EEG measures.

**Figure 1.**
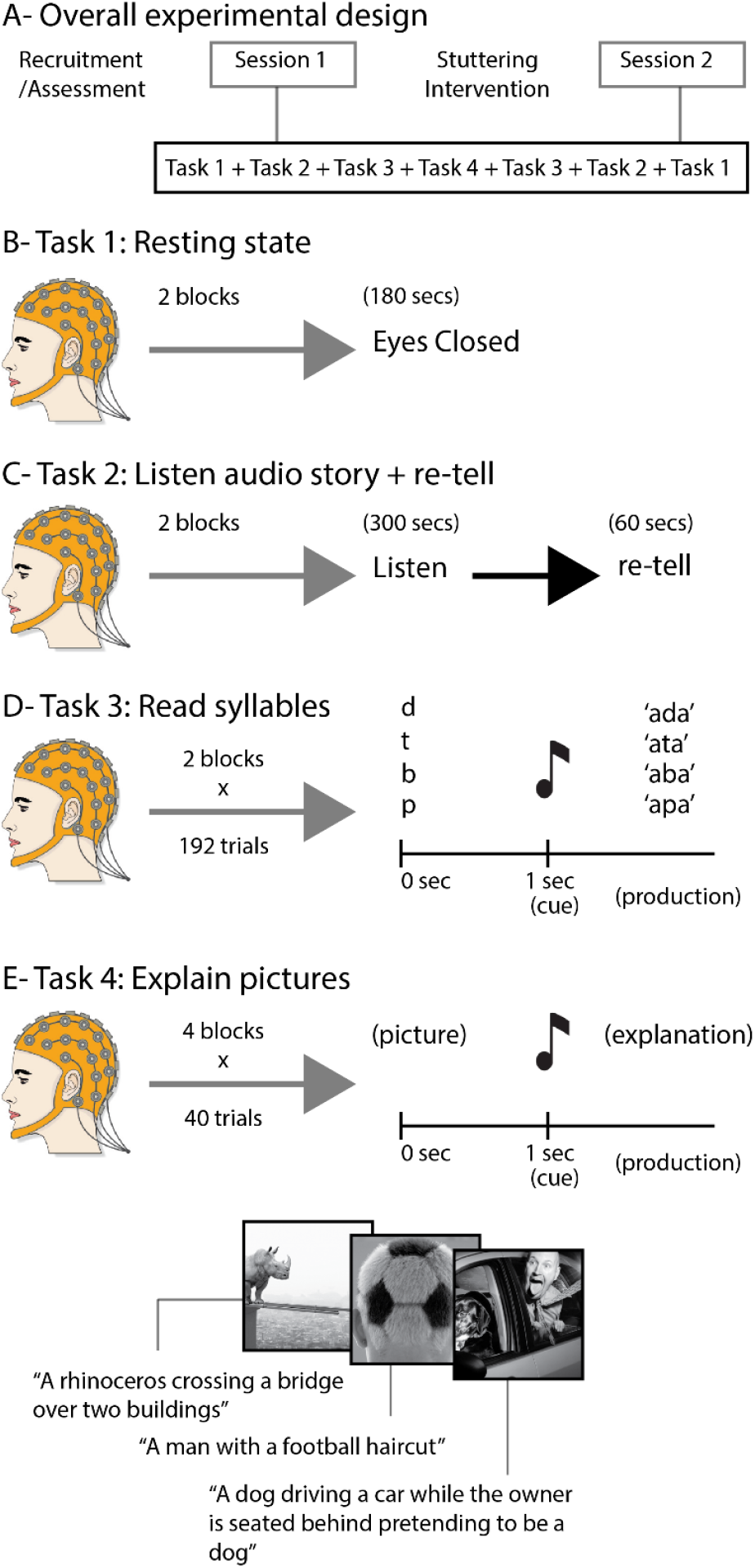
A-Two experimental sessions, one before and one after the stuttering intervention program. Within each session, 4 tasks were administered with EEG. B-Task 1 comprises a 3-minute resting state with eyes closed. C-Task 2 comprises listening and retelling an audio story. The perception component was 5 - minute long, and the retell limit time was 1 minute. D-Task 3 comprises reading syllables after an auditory cue (tone). Each block had a 7-minute duration. E-Task 4 comprises explaining pictures after an auditory cue. 4 blocks of this task were administered. Each block had an approximate duration of 3-4 minutes.

## 2. Material and methods

### 2.1 Participants

Seventeen participants (3 females, 1 left-handed), native Portuguese speaking, and aged between 18 and 50 years old (mean=33,2, sd=9,2) were recruited to this study with assistance from the Portuguese Stuttering Association (https://www.gaguez-apg.com) and local advertising among the University of Algarve (UAlg), Portugal (https://www.ualg.pt/). Participation was voluntary and participants gave their informed consent prior to the study and on each session. The study and experiments were conducted in accordance with the Declaration of Helsinki and Oviedo convention and approved by the ethical committee of UAlg. Participants were informed of the scope of the project, and the importance of taking part in both EEG testing phases (before and after speech therapy), as well as in all therapeutic sessions. One participant was excluded from group analyses due to dropout during the therapeutic phase. Three participants were excluded from group analyses due to technical issues in EEG recording and excess of movement artifacts. We note that the group sample had an unbalanced gender distribution, which should be considered when comparing the results of this study to other studies. For all speech conditions (i.e., overt speech production), participants were instructed to speak at a comfortable volume intensity, equivalent to a conversation with a friend standing one meter away. All participants reported normal hearing abilities and no history of psychiatric, neurological or language-related disorders apart from stuttering. This study was developed during the COVID-19 pandemic, a period with strong government (Portugal) restrictions to mobility and presential meetings, which limited the logistics of the study.

### 2.2 Intervention

Participants in this study received a total of 16 speech therapy sessions for stuttering, including 12 intervention sessions, 2 initial assessment sessions, and 2 final reassessment sessions. Each participant underwent therapy sessions over a period of 5 to 7 months. The sessions were scheduled weekly or biweekly, with an intensive mode employed to ensure effective learning (Irani *et al*., 2012). Therapeutic breaks were interspersed throughout the period to prevent therapist dependence and to promote the consolidation of the strategies and recommendations provided. These breaks were strategically planned and included home tasks to reinforce learning and helping participants to internalize the techniques more effectively. Participant recruitment and confirmation of the necessary inclusion criteria for this project were conducted during the initial assessment session.

The sessions were conducted online due to the COVID-19 pandemic restrictions in place. This format not only ensured the safety of all involved but also allowed to reach participants who were geographically distant. Several studies in the field of speech therapy provide evidence supporting the flexibility, effectiveness, and adequacy of online speech therapy (O’Brian, Packman and Onslow, 2008; Bridgman *et al*., 2016). The therapy was administered by two speech therapists with specialization in fluency disorders and extensive clinical experience, each with over 10 years of practice.

An integrative method was used that combined components of Stuttering Modification Therapy (Van Riper, 1973), Solution-Focused Brief Therapy (Burns, no date; McNeill, 2013), and Cognitive Behavioral Therapy (CBT) (Fry, 2013; Klein and Amster, 2018). As suggested by various authors, many individuals who stutter benefit from a therapeutic intervention that incorporates a blend of behavioral and emotional/cognitive-based approaches (Langevin *et al*., 2010; Beilby, Byrnes and Yaruss, 2012; Menzies *et al*., 2019).

Stuttering diagnosis and classification were confirmed through the analysis of speech samples obtained from: i) speech induced by responses to questions about films, hobbies, interests, and personal data; ii) description of images from the Frog Story book (Mayer, 1969); iii) description of images related to daily life themes; iv) reading passages from “The North Wind and the Sun” (Jesus, Valente and Hall, 2015) or “Arthur the rat” (Guimarães, 2002), according to individual reading habits. The analysis of the samples was conducted by our experts who classified stuttering using scales and a table of disfluency types.

Personal data collection, as well as information about clinical history, feelings, and thoughts over time related to coping with stuttering, and objectives and expectations regarding the therapeutic program, were also gathered. Additionally, the Portuguese version of the instruments OASES-A (Yaruss and Quesal, 2006), UTBAS (St Clare *et al*., 2009), and 4S (Boyle, 2013) were used to analyze participants’ communication, stuttering and speech self-perception using the Portuguese versions (UTBAS and 4S used translated versions developed by our team, *unpublished work in preparation*).

### 2.3 Assessment of self-perception and stuttering impact (OASES-A, 4S, UTBAS)

The three different questionnaires help to tailor therapy to individual needs by emphasizing areas that require most attention, including daily life challenges, stigma of being a PWS, and cognitive strategies. This approach fosters a more holistic treatment method. The three questionnaires were distributed in a randomized manner to ensure that participant’s responses reflected their immediate and current perceptions of stuttering. Participants were asked to respond to each questionnaire based on how they viewed and experienced their stuttering at that specific moment. This approach was chosen to capture the most accurate and timely insights into their individual experiences, reducing any potential bias that might arise from a fixed order of completion. By randomizing the order in which the questionnaires were presented, researchers aimed to obtain a more diverse and comprehensive understanding of the participants’ attitudes and feelings toward their stuttering, thereby enhancing the reliability and depth of the collected data.

OASES-A is a comprehensive tool that measures the impact of stuttering on a person’s life across multiple dimensions, including communicative competence and confidence, the speaker’s reactions to stuttering, and the functional and social limitations experienced due to stuttering. It provides clinicians and researchers with a detailed view of how stuttering affects an individual’s daily life and self-perception (Yaruss and Quesal, 2006, 2016). It consists of 100 items, each rated on a Likert scale from 1 to 5. It is organized into four sections, each corresponding to specific aspects of the World Health Organization’s International Classification of Functioning, Disability, and Health (ICF; World Health Organization [WHO], 2001): a) Section I -General Information: 20 items assessing how the speaker perceives their fluency and speech naturalness; b) Section II - Reactions to Stuttering: 30 items evaluating affective, behavioral, and cognitive responses to stuttering; c) Section III - Communication in Daily Situations: 25 items measuring difficulties in communication across various settings, including work, social interactions, and home; d) Section IV - Quality of Life: 25 items focusing on satisfaction with communication ability, personal and professional relationships, and overall well-being.

4S (Boyle, 2013) is a tool designed to measure the extent of self-stigma that individuals who stutter experience. It evaluates how deeply they internalize societal stereotypes and negative perceptions about stuttering. The 4S questionnaire comprises 6 blocks of questions with a total of 33 items. The 33 items of the 4S underwent factor analysis, and a three-factor solution was found to be the most parsimonious. The 4S measures awareness (e.g., “Most people in the general public believe that PWS are insecure”), agreement (e.g., “I believe that PWS are generally nervous”), and application (e.g., “Because I stutter, I feel less sociable than people who do not stutter”). The application section also includes items that measure behavioural outcomes resulting from self-stigma (e.g., “Because I stutter, I stop myself from taking jobs that require lots of talking”).

UTBAS (St Clare *et al*., 2009) (questionnaire facilitates the collection of information on the cognitive dimensions associated with stuttering. This scale identifies negative thoughts and beliefs that individuals who stutter might hold regarding their speech and the social repercussions of stuttering. The UTBAS scales offer clinicians a detailed checklist of negative thoughts experienced by adults who stutter. Developed through analysis of thought diaries from adults undergoing treatment for social anxiety and related psychological issues, the UTBAS helps identify and target specific thoughts for intervention in speech and psychological therapies. The scale includes 66 items: 27 specifically related to stuttering (e.g., “People who stutter are boring”) and 39 not directly related to stuttering (e.g., “People will laugh at me”). For each item, participants rate their responses on a 5-point scale as follows: 1 = never, 2 = rarely, 3 = sometimes, 4 = often, 5 = always. Participants are asked to evaluate each statement based on the following three parameters: a) Frequency: “How often do you have these thoughts?”; b) Belief: “How strongly do you believe these thoughts?”; c) Anxiety: “How anxious do these thoughts make you feel?”.

### 2.4 Experimental stimuli

In the ‘resting state’ task, a fixation-cross was used during the duration of the resting period (despite this task being done with eyes closed) and an auditory tone (800 Hz) with 50 ms duration (onset and offset ramp of 10 ms) indicated the end of the resting period (i.e., 180 sec). In the ‘listen-audio-story’ task, audio stories comprised short stories from native Portuguese speakers, each with 5-minute duration. In the ‘retell’ task, a fixation-cross was present throughout the period of the task. In the ‘read syllables’ task, written letters (‘B’, ‘D’, ‘P’, ‘T’) in upper-case and font size 24 were used. In the ‘explain pictures’ task, freely available photographs (https://unsplash.com/) were re-sized to 720 by 720 pixels and transformed into black & white. In the ‘read syllables’ and ‘explain pictures’ tasks, tones (400 and 800 Hz) were used to cue participants to initiate speech production on each trial. Auditory stimuli were presented binaurally at a comfortable intensity level. All text was displayed in Arial font-style and font-size 24 at the center of a gray screen using the ‘Psychotoolbox’ (in Matlab 2019).

Speech was recorded using a SM58-Shure (Chicago, USA) microphone, placed 20 cm away from the participants’ mouth. The microphone audio signal was integrated with the EEG signal using the Biosemi ERGO input, at a sampling rate of 16384 Hz (Biosemi, Amsterdam, The Netherlands).

### 2.5 Experimental procedure

The experiment was conducted in a sound-attenuated room. Participants were comfortably seated 50 cm from a screen and 20 cm from the microphone, and the quality of electrode impedance was kept below 10 kΩ.

The study consisted of two experimental sessions, one administered before and one after a speech therapy intervention. Each session was composed of four tasks (see figure 1) presented twice using a pyramid sequence presentation scheme. Task 1 (‘resting-state’) was presented at the beginning and end of the session, task 2 (‘listen and retell audio story’) was presented second from the beginning and second from the end, task 3 (‘read syllables’) was presented third from the beginning and third from the end, and task 4 (‘explain pictures’) was presented in the middle of the session split over four consecutive blocks.

‘Resting-state’ was performed with eyes closed during 3 minutes; three consecutive auditory beeps signalled the end of the resting block. ‘Listen and retell audio story’ comprised listening to a 5-minute audio story, following a 1-minute period during which participants were asked to provide an oral summary of the story. ‘Read syllables’ task involved a written consonant (i.e., the letter ‘B, ‘D’, ‘P’ or ‘T’) that participants were instructed to read aloud following the presentation of an auditory tone, 2 seconds after. To avoid articulatory preparation of the target consonant, participants were asked to introduce the vowel ‘a’ before and after the consonant, forming the respective items ‘aba’, ‘ada’, ‘apa’ and ‘ata’. The cue tone varied in frequency (400 Hz or 800 Hz) in a randomized order, with no implication on the task. The following trial was presented between 2 and 3 seconds (jittered) after the auditory cue. 192 trials were presented per block (7 minutes duration, 2 blocks per session). The ‘Explain pictures’ task followed the same presentation scheme as the read syllables task, but instead of a written letter, a black and white picture was presented. Following the auditory cue, participants were asked to describe aloud the corresponding picture. Once participants finished explaining a given picture, they pressed the ‘space bar’ of the keyboard to continue for the next trial. This task was divided in four blocks of 40 trials each. Between tasks, participants were given the opportunity to rest and drink water.

### 2.6 EEG data acquisition and preprocessing

Continuous EEG recordings were performed using the ‘BioSemi ActiveTwo’ 32-channel Ag/AgCl electrodes (10/20 system). The signal amplification was performed by the ‘BioSemi ActiveTwo’ amplifier, with a sampling rate of 16384 Hz during the entire period of the experiment and recorded by the ‘ActiveView600’ software (BioSemi, Amsterdam, The Netherlands). Conductive gel was applied using a plastic syringe. The 32 electrodes were then placed in the predefined sites: Fp1/Fp2, F3/F4, F7/F8, FT9/FT10, FC1/FC2, FC5/FC6, C3/C4, T7/T8, CP1/CP2, CP5/ CP6, TP9/TP10, P3/P4, P7/P8, O1/O2, Fz, Cz, Pz and Oz.

For all pre-processing stages and statistical analysis of the epoched EEG data, EEGLab (Delorme and Makeig, 2004) (version 2021) and Matlab (version R2022b) custom-made scripts were used. EEG data processing started with re-referencing the EEG channels to the Fz electrode, followed by band-pass filtering (0,5-40 Hz). Removal of signal artifacts was performed, and data were corrected for stereotypical artifacts related to eye movements, eye-blinks and/or noisy electrodes using independent component analysis (ICA, INFOMAX-runica algorithm as implemented in EEG Lab). Participant-specific independent components were categorized as neural activity or non-neural artifacts by visual inspection of their scalp topography, spectral peaks, and voltage range. Independent components labelled as non-neural artifacts were set to zero and the data matrix remixed accordingly to the EEGLab ICA procedure.

### 2.7 Data analysis

Two types of data analysis were performed in this study. The ERP analysis focused on the syllable reading and the picture description tasks. Both these tasks included a time-locking event (an auditory cue). The EEG classification of auditory cues focused on discriminating the identity of individual auditory cues based on their acoustic frequency (400 or 800 Hz) using multivariate pattern analysis.

#### 2.7.1 ERP analysis

An event-related potential (ERP) analysis was conducted for the ‘read syllables’ and ‘explain pictures’ tasks using the onset of the auditory cue as time-locking event (2000 ms after trial onset). For each trial, epochs of 500 ms and a 100 ms pre-stimulus baseline were referenced to the average of the baseline per EEG channel. Individual-subject ERPs obtained from the average of all trials were used to compute grand-average ERPs and group-statistics. The Cz channel was used for plotting the grand averaged ERP curves and performing the statistical analyses. A Time-window of interest was used to assess mean ERP responses per condition (i.e., P200 = [210-230 ms], as reported in the literature and showing salient results from the epoch analysis). Parametric paired t-tests were used to obtain values for statistical deviation from baseline and statistical comparisons between conditions. Reported statistical significance is p<0.05 for the windows-of-interest summaries and FDR corrected (q<0.05) for the continuous ERP analyses. Student t-tests (paired) were used to compare the individual ERPs to baseline averages, and to compare ERP results between sessions. Trials whose speech response onsets were shorter than 300 ms or longer than 1000 ms were removed from the analysis. Voice onset was computed using Matlab chronset_v1 toolbox.

#### 2.7.2 EEG classification of auditory cues

To investigate the specificity by which auditory cues are processed by the participants during speech preparation, we employed a binary classification of auditory cues. Cues were labelled according to their acoustic frequency (i.e., 400 or 800 Hz). Multivariate EEG signals (31 channels and 10 time-points) composed a pattern of EEG responses that was used to discriminate tones. Participants’ task involved waiting for the auditory cue to produce speech. Nevertheless, the frequency of the auditory cue is irrelevant to the task. Tone frequency discrimination may require auditory processing that competes with that necessary for speech prediction and monitoring. Hence, it is likely that EEG-based tone discrimination may be related to fluency. Multivariate classification was employed to investigate whether specific temporal EEG signatures enable tonal discrimination. To this end we used a supervised machine learning algorithm (linear support vector machines, linear-SVM; Cortes and Vapnik, 1995) as implemented in the Bioinformatics Matlab toolbox using the SMO algorithm. We employed a linear kernel for a more direct interpretation of the classification weights obtained during training. SVM regularization was further used to account for MVPA feature outliers during training, which would otherwise risk overfitting the classification model and reduce model generalization (i.e., produce low classification of the testing set). Regularization in the SMO algorithm is operationalized by the Karush-Kuhn-Tucker (KKT) conditions. Binary classification produces an accuracy, for which 0.5 (i.e., 50%) is the chance level. Accuracies significantly above this value are informative of tonal discrimination in the EEG trial-by-trial responses. For plotting the accuracies in figure 3C, we chose to subtract the chance-level (0.5) value from the accuracy results. For this reason, our accuracy scores are centred around zero, with values above zero representing tonal discrimination.

Cross-validation of the multivariate classification analysis is necessary, which is accomplished by splitting the data into training and testing datasets. We used a cross-validation approach that split the data in 10 equal parts, using 9 parts for training and 1 part for testing, and repeating the training-testing partitioning 10 times. Each partition of the data was used only once as a testing set. Cross-validation assures that training and testing sets are independently processed.

## Results

Speech was assessed based on dysfluency scores in two separately tasks: the story retell task and the picture description task. In the story retell task, four additional variables were obtained: the dysfluency duration, the tension degree, the existence of secondary or concomitant stuttering behaviours, and the impoverished naturalness of speech. Fluency scores were obtained before and after stuttering intervention. Overall, stuttering intervention reduced the dysfluency scores obtained in both tasks. This reduction was significant in the story retell task (t=2.69(12), p=0.019), but not in the picture description task (t=1.73(12), p=0.11) (see figure 2A). An interesting observation in the picture description task was that after speech therapy intervention, participants tended to describe the images in more detail, which suggests a reduction of their avoidance behaviours. Despite this fact, PWS showed a reduction of dysfluency rate after intervention.

**Figure 2.**
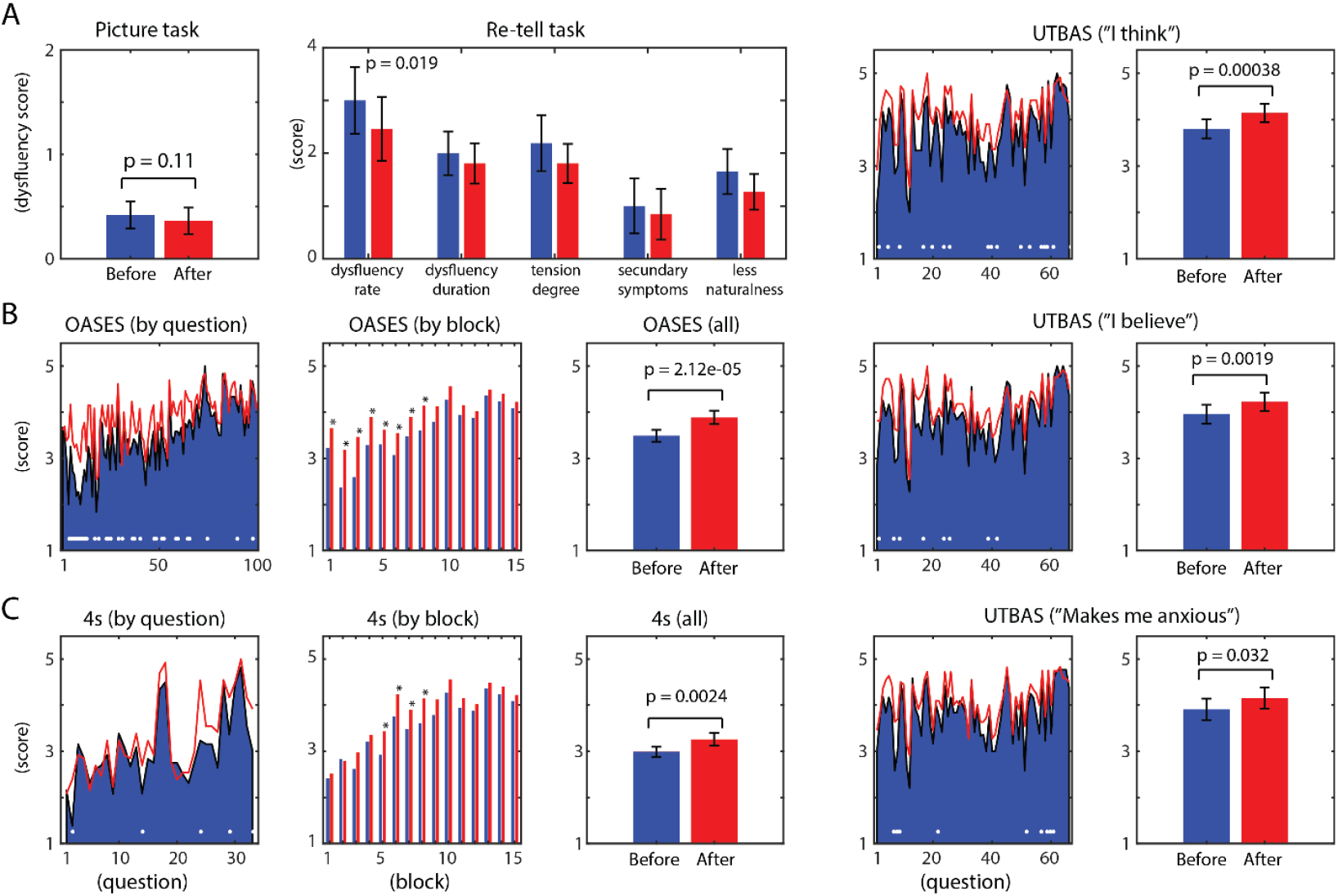
Behavioral results. Before speech therapy in blue and after in red. A-Dysfluency scores obtained from the picture description task (no stuttering = 0, stuttering = 1, severe stuttering = 2) and from the retell audio story task (scale 0 to 5 of dysfluency rate, duration, tension, secondary symptoms and lack of naturalness). B-Results from the OASES-A questionnaire. Left: results from all individual questions (white dots below depict statistical differences, p<0.05). Middl e: results averaged by block (asteriscs over the bar pairs depict statistical differences, p<0.05). Right: results from all questions combined. C-Results from the 4s questionnaire. Left, middle and right plots depict the results by question, by block and combining all questions, respectively. D-Results from the UTBAS questionnaire. Left and right plots depict results by question, and combining all questions, respectively. Top, middle and bottom plots reflect the 3 dimensions of the UTBAS questionnaire: “I think”, “I believe”, and “makes me anxious”, regarding each statement of the questionnaire.

The OASES-A questionnaire was administered before and after stuttering intervention, with an overall improvement of results (see figure 2B). Note that OASES-A measures were adapted such that higher scores represent positive indicators. This allowed us to group the OASES-A scores by section, as well as in a single summary value. We found significant positive changes in several individual questions, and by group in the first 8 sections of the OASES-A questionnaire, showing that participants change the perception of their communicative skills towards a more positive view (p<0.05). Overall, the combination of all questions into a single value improved significantly with therapy (t=-7.05(11), p=2.12 e-5).

The 4S questionnaire also allowed for the analysis of self-perception related to stuttering, providing insights into how participants internalized negative perceptions and societal stereotypes about stuttering evolved in result of the intervention. Analysis of the results revealed statistically significant changes in self-stigma levels. Specifically, participants showed a significant reduction of self-stigma in blocks 5, 6, 7, and 8 (p<0.05), while the global 4S summary revealed significant (t=-3.83(12), p=0.0024).

Finally, the UTBAS questionnaire revealed significant differences across the three evaluated parameters. Specifically, the results indicated that after therapy, participants reported a lower frequency of having the thoughts measured by the questionnaire (t = -5.03, p = 3.85e-04). The strength of belief in these thoughts was also significantly reduced (t = -4.05, p = 0.0019). The level of anxiety induced by these thoughts showed a smaller, yet still significant effect (t = -2.46, p = 0.032).

Electrophysiological changes were observed in our task, as related to event related potentials time-locked to the auditory cues. Grand-average ERPs produced the expected N100 and P200 peaks, both for the syllable task (figure 3A) and the picture description task (figure 3B), corrected for multiple comparisons (i.e., multiple time-points) using FDR correction. Significant changes in ERPs across the epoch window were present (p<0.05, uncorrected) mainly for the P200 peak in both ERP production tasks (i.e., syllables and picture tasks). A summary of the P200 peak (i.e., averaged time-points between 2180-2220 ms, which are 180-220 ms after the auditory cues) was used to assess possible statistical differences between sessions, and to compute Pearson correlations to the dysfluency scores obtained from the story retell task. In both tasks, there was a trend for reducing the magnitude of the P200 peak from pre-therapy to post-therapy, though not statistically significant (t=1.78(12), p=0.099 for the syllable task, and t=1.66(12), p=0.12 for the picture task).

**Figure 3.**
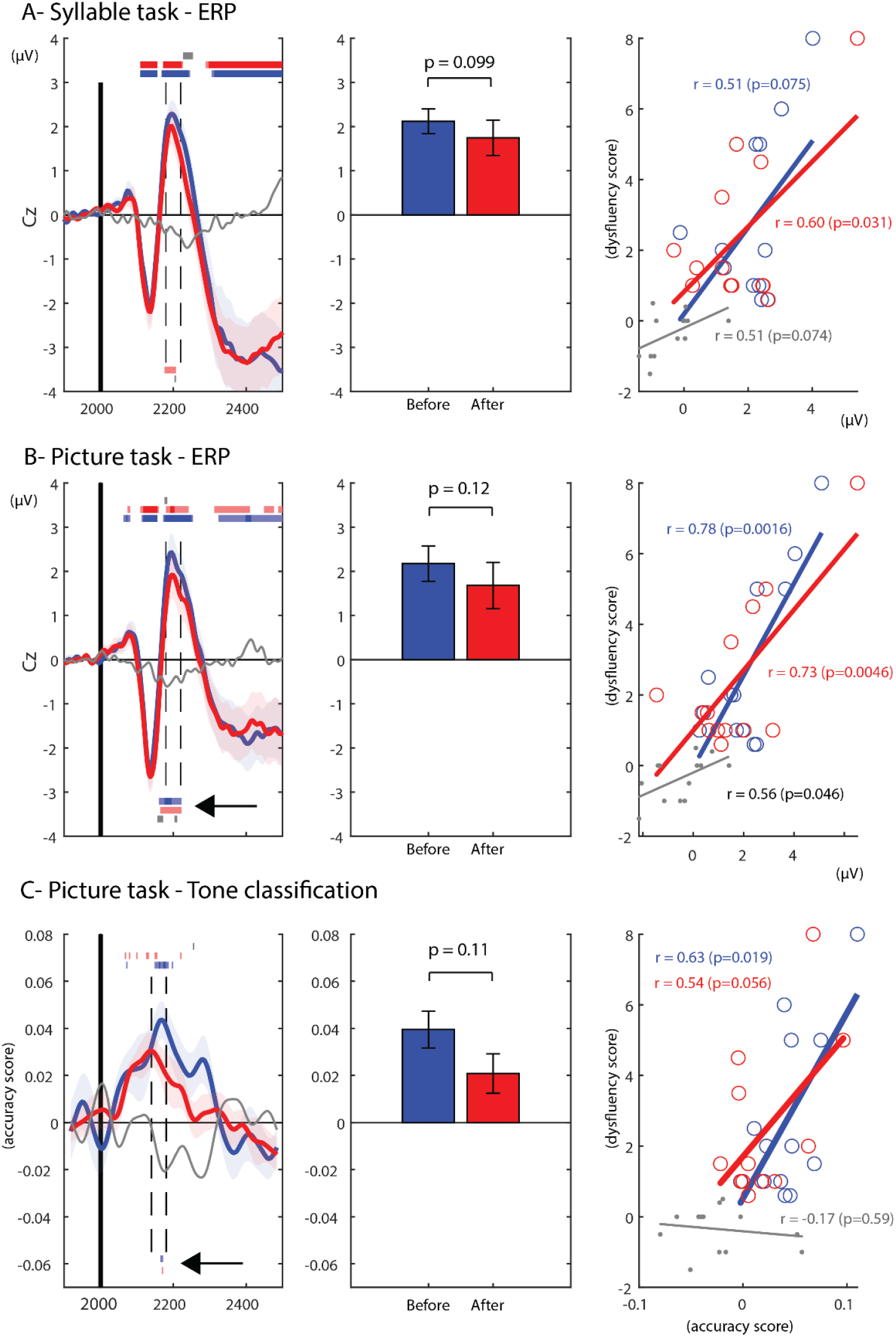
EEG auditory responses. A-Syllable task. Left: grand-average ERP curves time-locked to the onset of the auditory cue. Red and blue shadings along the curves reflect standard error of the mean (std); blue is before and red is after speech therapy; gray thin line depicts difference (after-before); horizontal bars depict statistical significance (p<0.05) in light shades, and corrected p-values (FDR, q<0.05) in dark shades; upper horizontal statistical lines reflect deviations from baseline; lower horizontal statistical lines reflect correlation to fluency across participants; gray bars depict statistical differences between sessions. Middle: bar-graph of ERP summaries from the window of interest centred in 2180-2220 ms (180-220 ms after auditory cue). Right: scatter plot of all participants depicting the relation between P200 averages and speech fluency. Gray color depicts P200 across dysfluency across sessions (after-before); red, blue and gray lines are best fit lines. B-Picture description task. Same organization as in the syllable task. C-Tone discrimination results (multivariate classification of individual tones -400 vs. 800 Hz) during the picture description task. Black arrow in the left plot highlights significant correlations (uncorrected) between tone discrimination and fluency, both before and after intervention, in overlapping time windows.

Pearson correlations between the P200 peak summaries and the dysfluency scores were considerable in both tasks. For the syllable task, we observed a non-significant correlation of r=0.51 (p=0.075) before speech therapy and a significant correlation of r=0.60 (p=0.031) after speech therapy, as well as a non-significant trend for the correlation of the ERP differences with the improvement of fluency between sessions (r=0.51, p=0.074). Importantly, in the more ecological speech task of picture description, Pearson correlations between the P200 peak summaries and the dysfluency scores were significant. We observed positive correlations before speech therapy (r=0.78, p=0.016]) and after speech therapy (r=0.73, p=0046). Interestingly, and although marginal, we found that improvements in speech fluency (i.e., reductions in dysfluency scores) correlated with reductions in the P200 potentials across our group of participants (r=0.56, p=0.046).

To further assess whether auditory attention has an influence in stuttering, we employed multivariate classification of the two tones used as speech production cues (i.e., 400 Hz versus 800 Hz tones). We found that tonal discrimination was possible based on EEG response patterns obtained between 2140-2180 ms (i.e., between 140-180 ms after the auditory cue), with averaged classification accuracies reaching 0.54 (see figure 3C, note that for plotting, accuracies were transformed to accuracy scores by subtracting chance-level values – 0.5). Despite pre-therapy showing higher accuracy scores compared to post-therapy, such difference was not statistically significant (t=1.73(12), p=0.11). Furthermore, tone discrimination was significantly correlated with dysfluency scores before (r=0.63, p=0.019), but not after speech therapy (r=0.54, p=0.056), despite the statistical trend.

## 4. Discussion

Speech production requires fine-tuned sensorimotor circuits able to link speaking intentions to sensory predictions and the subsequent processing of sensory consequences of speech (Tremblay, Deschamps and Gracco, 2016). From speech development to everyday speech control, the association between motor speech planning and speech perception may constitute a central mechanism by which humans master to learn and produce fluent speech (Hickok, Houde and Rong, 2011; Poeppel *et al*., 2012). Exposure to the auditory consequences of self-produced speech may help training the sensorimotor network during childhood (Chang *et al*., 2019). Despite stuttering in childhood having a considerable prevalence (5-10 %), the recovery rate is exceptionally high (80%), which may reflect sensorimotor plasticity at young age (typically below 5 years old) (Chang and Zhu, 2013; Yairi and Ambrose, 2013). In contrast, when stuttering persists to adulthood, it is highly resistant to treatment. Overall, our ability to address stuttering in adults and ultimately develop effective therapeutic approaches depends on our knowledge of the brain dynamics responsible for speech production in PWS, and their relation to individual stuttering variability. Here, we unveil a relationship between stuttering variability and EEG responses to auditory cues, while PWS prepared to speak. The modulation of auditory brain responses by stuttering severity may be interpreted as contributing for stuttering (Civier, Tasko and Guenther, 2010), as a compensatory mechanism to deal with insufficiently activated parts of a broader speech production circuitry (Neef *et al*., 2015), or as a bi-product of stuttering with no causal relationship.

We intended to study the involvement of the auditory cortex in speech preparation by probing participants with auditory stimuli during speech preparation. Although indirect, the assumption that an EEG response to external auditory probes may reflect speech prediction efforts, has been previously suggested (Daliri 2015, 2018). Here, we adopted a compatible strategy by employing auditory cues for speech production. In previous research (Daliri 2015), auditory probes across trials were presented only in a subset of trials, and at jittered times (i.e., unpredictable to the participants). In our case, auditory probes were presented exactly 2 seconds after the target visual stimuli and used to cue participants to initiate their vocal response. Importantly, the presentation of the auditory probes is expected to compete for neural resources related to sensorimotor aspects of speech production, namely the auditory prediction and preparation for speech monitoring. It is possible that different experimental details can affect the EEG responses underlying speech preparation. Our decision to employ an auditory probe that functioned simultaneously as the cue for production relied on its potential to convey an intuitive task that would not attract undesirable (or inconsistent) attention to the auditory probes. The fact that we observe a link between the level of EEG response to the auditory cues and stuttering is informative for the potential of this paradigm to tap onto the sensorimotor basis of speech production and of stuttering.

Alternatively, it is possible that our task is related to rhythm, which is a known phenomenon influencing fluency in stuttering (Sares *et al*., 2019), as well as motor control (e.g., in PD) (Bella *et al*., 2015). The auditory cue was presented at fixed times after the target visual stimulus (2 seconds after). Consistent inter-stimulus times offered predictability in our task, which was designed to reduce auditory attention and make the task as natural and fluid as possible. Increased EEG responses to the auditory cues in participants with more severe stuttering may reflect lack of endogenous rhythm capacity, and in turn over-dependency from the exogenous (i.e., external) auditory cues, leading to the exacerbated neural responses (Jenson *et al*., 2020). Rhythm in motor control depends on cortico-basal-ganglia-thalamo-cortical circuits, likely involving the hyperdirect pathway (HDP), which has been more recently implicated in stuttering (Usler, 2022). Although the effect of rhythm is interesting as a potential alternative explanation to our task, future experimental protocols, for example contrasting auditory versus visual cues, as well as jittered versus fixed cue presentation times may be relevant strategies to further explore this possibility.

In addition, to provide support for an over activation of the auditory cortex during speech preparation in more severe stuttering participants, we assessed auditory specificity based on tonal discrimination. Tones consisted of 400 and 800 Hz sinusoidal signals, presented randomly with equal probabilities. Participants were not informed of the relevance of this stimuli variability. By exploiting the benefit of multivariate pattern analysis applied to EEG (Hausfeld *et al*., 2012; Correia *et al*., 2015), we assessed the degree of tonal discrimination for each participant and session throughout the time-course of our epoch. Despite participants have received no indication for the importance of tonal discrimination, their EEG responses conveyed such information, particularly in the time window overlapping with the P200 potential. This finding was expected based on previous studies showing task-relevant and task-irrelevant auditory stimuli discrimination (Hausfeld 2011). Interestingly, the degree of tonal discrimination was positively correlated to dysfluency scores. This novel result indicates that participants with more severe stuttering are also more attentive to the content of the external stimuli. Overall, our findings suggest that discrimination analyses using multivariate pattern analysis can provide valuable indications for the level of processing that stuttering affects participants. Future EEG studies capitalizing on multivariate pattern analysis may uncover several hidden aspects of the neural mechanisms of stuttering, including auditory discrimination, articulatory discrimination, or voicing discrimination across brain regions and time-windows of interest (Cheung *et al*., 2016; Correia *et al*., 2020).

Stuttering interventions have evolved considerably in the past decades (Laiho *et al*., 2022). Speech therapy treatments intended to modify the speaking technique are changing towards holistic strategies that contemplate factors typically pertained as secondary, such as speaking confidence or speaking diversity (Hart, Breen and Beilby, 2021; Lamoureux *et al*., 2024). Fluency improvement obtained from these approaches are likely the indirect consequence of speech therapy instead of a fixed or learned behaviour. Beyond fluency, we assessed self-perceptions of a group of PWS before and after an intervention program focused on various aspects of communication and psychological well-being. The intervention aimed to enhance overall communication effectiveness and address related emotional and cognitive factors. Our results suggest, indeed, that the variability in stuttering may be more reflective of the stuttering state rather than a stable trait. It is important to note that this variability is a constant factor across all participants. Despite the inherent variability typically observed at the individual level, we are still able to identify significant differences between sessions. This indicates that, even with the expected fluctuations in stuttering severity, meaningful changes were detected, highlighting the sensitivity and robustness of our approach in capturing the effects of the intervention.

Finally, by including a flexible speech production task (i.e., the picture describing task), which demanded participants to plan and encode their vocal responses (i.e., without explicit speech targets, such as in reading, naming or repetition), we intended to investigate the neural basis of stuttering under more natural/ecological conditions. Beyond the analyses reported, we expected participants to show inter-trial fluency variability that would enable employing classification of fluent vs. dysfluent trials within each participant. This aspect is in turn relevant to distinguish state from trait in stuttering (Belyk, Kraft and Brown, 2015), and more clearly study the EEG signatures of stuttering. However, our findings revealed insufficient balance in fluent/dysfluent trial ratios in the picture description task. Only two participants showed a balanced ratio of fluent/dysfluent trials, which were insufficient to investigate this dimension in our data. It is difficult to predict individual dysfluency, at the individual-trial level. A possible direction for future research may reside in evaluating EEG response dynamics in continuous speech production. Brain speech entrainment methods applied to continuous EEG data have been successful in unveiling the link between ongoing speech acoustics and EEG signals in speech perception (Peelle and Davis, 2012; Ghitza, Giraud and Poeppel, 2013). Similar methods have been used in intracranial EEG (Brumberg *et al*., 2016) during speech production, but their use in extracranial EEG has not been established.

## 5. Conclusions

Our findings suggest a positive correlation between stuttering severity and the modulation of auditory brain responses, indicating that individuals who stutter may exhibit increased auditory cortical activation during speech preparation. The use of auditory stimuli as cues for speech production was intended to investigate how these stimuli compete for neural resources involved in speech prediction and preparation. A more fine-grained analysis of the EEG responses to the auditory cues revealed that tonal discrimination was possible in our group of participants around the time window of the P200 ERP response, and in turn correlated to dysfluency rates, suggesting that participants with more severe stuttering are more attentive to external stimuli during speech preparation (Civier, Tasko and Guenther, 2010). Despite the inherent stuttering variability, significant differences were observed in self-perception and communication effectiveness before and after an intervention program. This highlights how effectively the approach can identify therapeutic outcomes that extend beyond gains in fluency. Future research should further explore how auditory discrimination and other cognitive factors are affected by stuttering using multivariate pattern analysis. Understanding these dynamics could help in developing more effective, holistic interventions that address not only fluency but also psychological and emotional aspects of stuttering.

Our data is available upon request.

## 6. Acknowledgements

We are grateful to the Portuguese association of persons who stutter (APG) for their help in the divulgation our study and recruitment of participants; to the clinic PIN, Portugal, for letting us use their facilities; and to the group of cognitive neurosciences (GNC), CUIP, University of Algarve, for continuous feedback and support throughout the project.

## 7. Author contributions

MR was involved in study conceptualization and writing. JC helped conceptualizing the study and participated in writing the manuscript. Both JC and MR were responsible for the stuttering intervention procedure development, recruitment of participants and evaluation of stuttering and fluency metrics. JMC conceptualized the EEG experimental paradigm, acquired and analysed the EEG data, and participated in writing the manuscript.

## 8. Funding

None of the following funding sources were involved in any stage of the present study.

*This work was supported by the Portuguese science organization (FCT: ‘Fundação para a Ciência e Tecnologia’) [grant number CEECIND/03696/2017]; the Portuguese Society for Speech Therapy (SPTF) in collaboration with Pharmis, Portugal; the University Hospital Centre of Algarve (CHUA), Portugal;* Alcoitão School of Health Sciences, Portugal; *and the municipality of São Brás de Alportel, Portugal*.

## Notes

### Competing Interest Statement

The authors have declared no competing interest.

